# Directional translocation resistance of Zika xrRNA

**DOI:** 10.1101/2020.06.17.157297

**Authors:** Antonio Suma, Lucia Coronel, Giovanni Bussi, Cristian Micheletti

## Abstract

xrRNAs from flaviviruses survive in host cells for their exceptional dichotomic response to the unfolding action of different enzymes. They can be unwound, and hence copied, by replicases, and yet can resist degradation by exonucleases. How the same stretch of xrRNA can encode such diverse responses is an open question. Here, by using atomistic models and translocation simulations, we uncover an elaborate and directional mechanism for how stress propagates when the two xrRNA ends, 5′ and 3′, are driven through a pore. Pulling the 3′ end, as done by replicases, elicits a progressive unfolding; pulling the 5′ end, as done by exonucleases, triggers a counterintuitive molecular tightening. Thus, in what appears to be a remarkable instance of intra-molecular tensegrity, the very pulling of the 5′ end is what boosts resistance to translocation and consequently to degradation. The uncovered mechanistic principle might be co-opted to design molecular meta-materials.

## INTRODUCTION

Subgenomic flavivirus RNAs (sfRNAs) are non-coding RNAs that result from the partial degradation of the viral genomic RNAs by cellular Xrn1 exoribonucleases. They accumulate in cells infected by flaviviruses, such as Zika, disrupting several molecular processes of the host. While sfRNAs cannot be degraded by exonucleases, they can still be processed by polymerases and reverse transcriptases [1–5].

As part of the ongoing endeavor to explain sfRNAs dichotomic response to processive enzymes, which is at the basis of flaviviruses pathogenicity, a series of targeted *in vitro* assays, including mutagenesis experiments have been carried out. They clarified that resistance to exonucleases is provided by a ca 71nucleotide-long sub-sequence, xrRNA in short, that can withstand degradation at its 5′ end by various other exonucleases besides Xrn1 [5]. Inspection of xrRNA structure —the one of Zika is shown in Fig. 1— reveals multiple pseudoknots and a ring-like motif that appears well-suited to dock onto the surface of the exonuclease [2, 3].

**FIG. 1.**
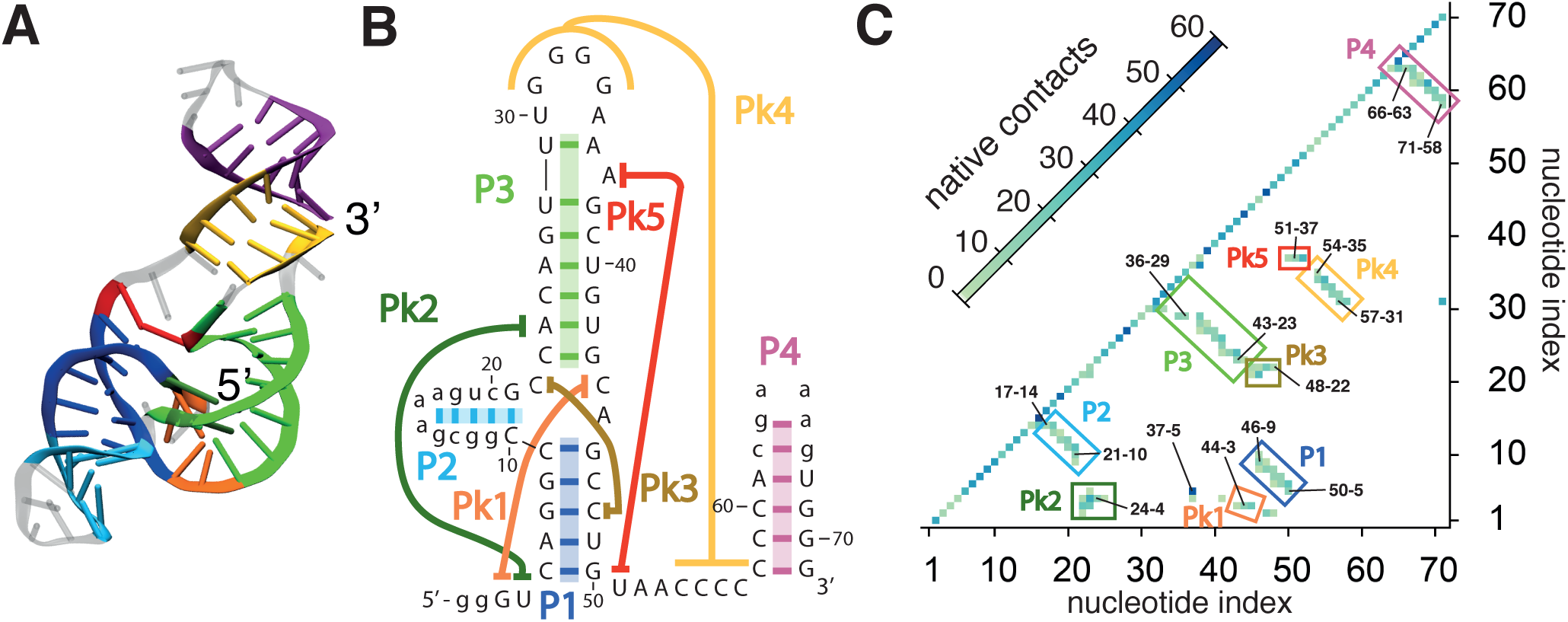
Structural organization of Zika xrRNA. (A) Ribbon representation of the 71-nucleotide long xrRNA of the Zika virus, PDB entry 5TPY [3]. (B) Diagrammatic representation of canonical and non-canonical base pairings, as annotated by Barnaba [17]. (C) Heat map of the native contacts. The map includes heavy-atom interactions within a 4Å cutoff distance and thus include contacts formed by bases, sugar, and phosphate moieties. The four helices, P1 to P4, and pseudoknotted regions, Pk1 to Pk5, are highlighted. Diagram in panel B follows ref. [3]. Secondary elements are highlighted with the same color code across the three panels. Structural renderings are based on the VMD software package [18].

The elaborate architecture of the relatively-short xrRNAs poses general questions about the compliance of xrRNAs to being translocated through the lumen of enzymes working from the 5′ to 3′ ends, as done by exonucleases, or from the 3′ to 5′ ones, as done by replicases and reverse transcriptases. What are the mechanistic and thermodynamic underpinnings of xrRNA resistance? Would the same directional resistance be observed in single-molecule setups forcing translocation through narrow pores? And, can one learn general tensegrity principles transferable to molecular design?

Prompted by these questions, we set out to study the microscopic mechanical underpinnings of sfRNAs resistance by using an atomistic model of the Zika xrRNA and stochastic simulations of its driven translocation through a narrow cylindrical pore. The pore-translocation setup provides a convenient abstraction of the resistance of sfRNAs to being driven through various types of exonucleases. In a similar spirit, our model system is exclusively informed with the native three-dimensional organization of the Zika xrRNA, thus discounting sequence-dependent effects and specific interactions with nucleases. More in general, it allows for a comparison with other biologically-motivated contexts where the intra-molecular organization has been investigated in connection with the compliance to translocate through enzymes or synthetic pores [6–16].

In our systematic study, we perform hundreds of force-ramped translocation simulations from the 3′ and 5 ends. Structural effectors of translocation compliance are established by analysing the network of strained interactions as well as by selectively removing groups of them. Bell-Evans analysis and especially metadynamics simulations are then used to obtain a characterization of the free-energy landscape, of the barriers associated with translocation resistance and of the pathway to the effective transition state.

Throughout the explored combination of ramping protocols and temperatures, we observe that translocation from the 5′ end requires forces that significantly exceed those needed for the 3′ end. This shows that Zika xrRNA architecture, which is the sole input of our model, suffices to encode a significant directional response to translocation. The enhanced hindrance at the 5′ end is shown to originate from an elaborate stress redistribution mechanism that has no analog at the 3′ end. The much higher free-energy barrier encountered at the 5′ end reflects an orders of magnitude difference of the time required to activate translocation from the two ends at constant force.

The results illuminate the structure-based contribution of xrRNA directional resistance and provide a single quantitative framework that recapitulates the resistance to 5′ → 3′ exonuclease degradation and 3′ → 5′ processability by reverse transcriptases and polymerases. It also indicates that pore-translocation setups, which could verify our simulation results, could be used as valuable probes of xrRNA resistance.

## RESULTS

### Structure of Zika xrRNA

Fig. 1A shows the structure of the Zika xrRNA, a 71-nucleotide long non-coding subgenomic RNA that can resists degradation by Xrn1 and other 5′ → 3′ ribonucleases [3]. Its canonical and non-canonical base pairings are summarised in Fig. 1B. Heavy atoms contacts, including those of sugar and phosphate moieties, are shown in the native contact map of Fig. 1C. The secondary and tertiary organization is articulated over four helices (P1 to P4), and multiple pseudoknot-like elements (Pk1 to Pk5).

### Response to pore translocation

We integrated atomistic modelling, molecular dynamics simulations, force-spectroscopy analysis and thermodynamic sampling techniques to understand how exactly the molecular architecture of the Zika xrRNA underpins its resistance to being driven through the lumen of exonucleases from the 5′ end, while still allowing to be processed and unwound by enzymes, such as polymerases and reverstranscriptases, which work from the 3′ end.

Inspired by force-spectroscopy approaches[19, 20], we adopted a force-ramping protocol where the Zika xrRNA is driven through a cylindrical pore. The pore is 11.7Åwide and runs perpendicularly through a parallelepiped slab that is 19.5Åthick, Fig. 2A. The pore geometry approximates the Xrn1 nuclease lumen [21] and allows the passage of only a single RNA strand at a time. Mimicking an electrophoretic setup [11, 22], the driving force was applied exclusively to the xrRNA *P* atoms inside the pore, whose longitudinal axis we take as the vertical or *z* axis of a Cartesian coordinate system, with the *x* − *y* plane coinciding with the *cis* surface of the slab.

**FIG. 2.**
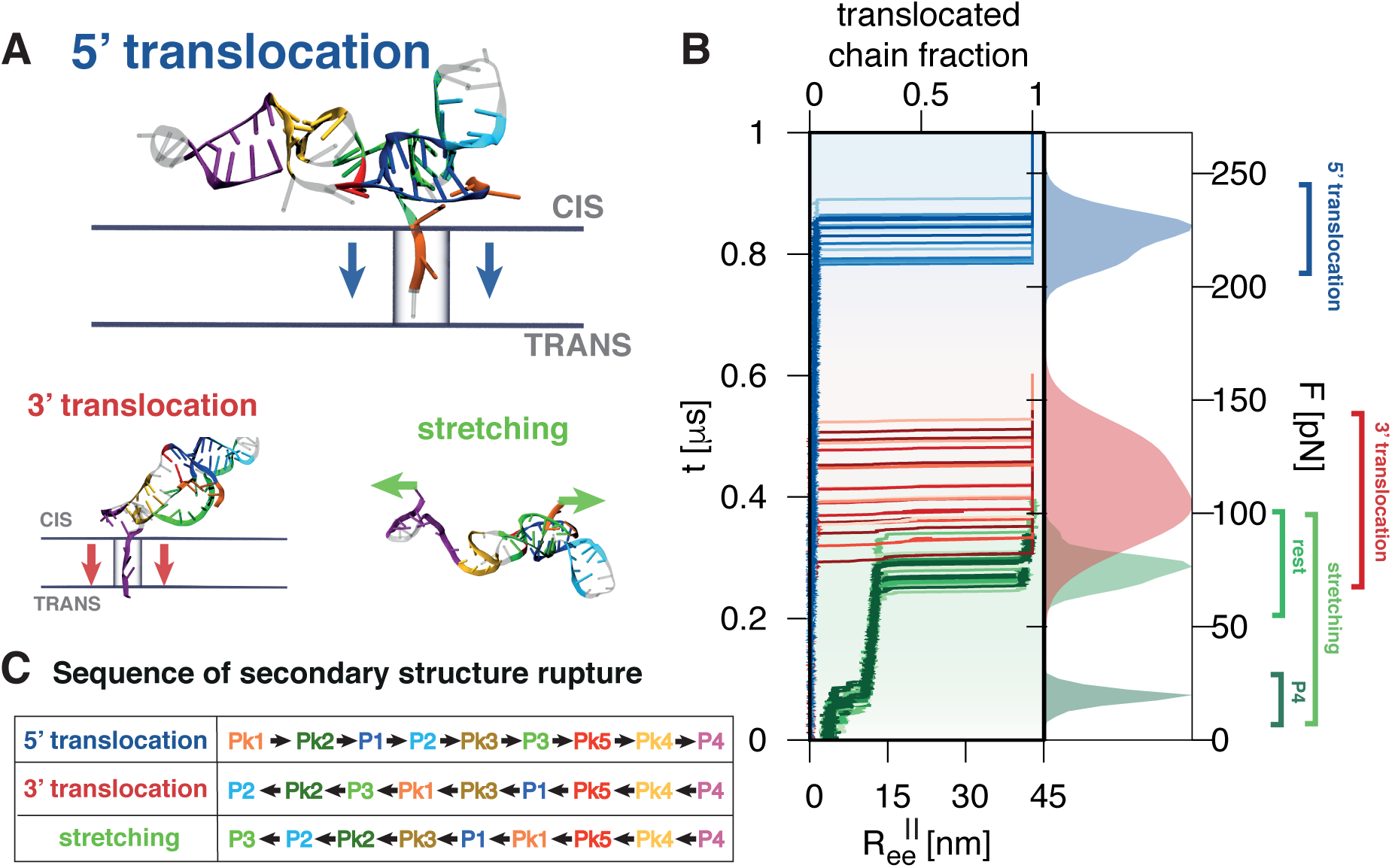
Zika xrRNA response to pore translocation and stretching. (A) Typical snapshots of the initial stages of translocation and stretching processes; the arrows indicate the direction of the force applied inside the pore or at the chain ends. (B) Temporal trace of the translocated chain fraction from the 5′ (blue) and 3′ (red) ends for 20 trajectories at the reference temperature *T* = 300*K* and force-ramping rate *r* = *r*_0_ = 270pN/*µ*s. The green traces show the longitudinal end-to-end distance, 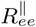 for 20 stretching trajectories at the same values of *T* and *r*. The filled curves on the right are the associated height-normalised distributions of the critical forces for translocation and mechanical unfolding, obtained with Gaussian kernel density estimators. (C) Typical order of rupture of native contacts during translocation and stretching, see also Fig. S2. Secondary elements in panels (A), (C) are highlighted with the same color code of Fig. 1A.

The xrRNA structure was represented with SMOG. The model retains the full atomistic details and has been validated for various molecular systems, including nucleic acids [23–26]. SMOG employs a native-centric force field that allows us to highlight structure-dependent properties while filtering out details more related to sequence or specific RNA-protein interactions.

We carried out hundreds of ramped translocation simulations of the molecule after priming it at the pore entrance from both ends, Fig. 2A. Fig. 2B shows a representative set of translocation trajectories from 5′ and 3′ ends. The curves show the temporal trace of the translocated fraction of the molecule at the reference temperature *T* = 300K and force-ramping rate *r*_0_ = 270pN/*µ*s. Translocation initiates only after a long loading stage, after which it proceeds precipitously. Major differences in translocation resistance emerge between the 5′ and 3′ ends.

The unlocking, or triggering of translocation, occurs when the vertical position of the leading *P* atom, measured relative to the *cis* plane, falls below *z* = − 19.5Å or *z* = − 15Å for 5′ or 3′ entries, respectively, and is followed by a cascade of ruptured contacts without significant pauses between the breaking of various secondary elements (Supplementary Figs. 2, 3). The typical order of rupture is given Fig. 2C. For the inherent stochasticity of the process, the triggering force is not unique but varies in a range, following a unimodal probability distribution as shown in Fig. 2B. Compared to the 3′ case, the most probable triggering force at the 5′ end is more than double; it is also much larger than the standard deviation of the triggering forces at the same end, 8pN, or at the opposite one, 17pN.

### Comparison with xrRNA stretching response

The unusually high resistance at the 5′ end is further highlighted by contrasting the translocation forces at this end with those required to unfold the xrRNA by pulling the two termini apart, as in force-spectroscopy. As shown in Fig. 2B, unfolding by stretching, which occurs when the longitudinal end-to-end distance, 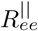 exceeds 16nm, requires forces three times smaller than those needed to initiate 5′ translocations. Interestingly, the typical sequence of secondary elements rupture by stretching is similar to the case of 3′ translocations. The main difference is that the P4 helix, which yields when 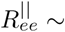 9nm, now unfolds well before the rest of the structure (Fig. 2B and Supplementary Fig. 2).

In summary, at similar conditions, the characteristic force needed to initiate xrRNA translocation from the 5′ end, which resists exonuclease degradation, is distinctly higher than the forces that causes unfolding by stretching or by translocation from the 3′, which is processable by replicases. The same results are seen for the several hundreds of trajectories gathered over 19 combinations of *T* and *r*, from lower to higher temperatures and faster or slower ramping protocols than the reference values, see Supplementary Figs. 4 and 7.

### Bell-Evans analysis

The extensive set of translocations at all explored (*T, r*) combinations can be recapitulated by the Bell-Evans (BE) analysis. In force-spectroscopy contexts it is customary to resort to the following BE expression for the most probable rupture force [19, 20, 27]:

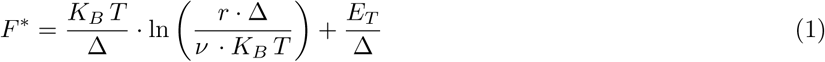

where *v* is a kinetic coefficient and *E*_*T*_ and Δ, the key quantities of interest, are the effective height and width of the thermodynamic barrier associated to the rupture event at zero force.

Fig. 3A shows that expression 1 provides a good fit of the most probable translocation forces, yielding a *χ*^2^ ∼ 3, hence of order unity, for both the 5′ and 3′ ends when all 19 (*T, r*) combinations are considered simultaneously. For clarity, only a subset of the considered datapoints are shown in Fig. 3 – see Supplementary Fig. 7 for the complete set – and the best fit to the BE expression is shown for *T* = 300K only.

**FIG. 3.**
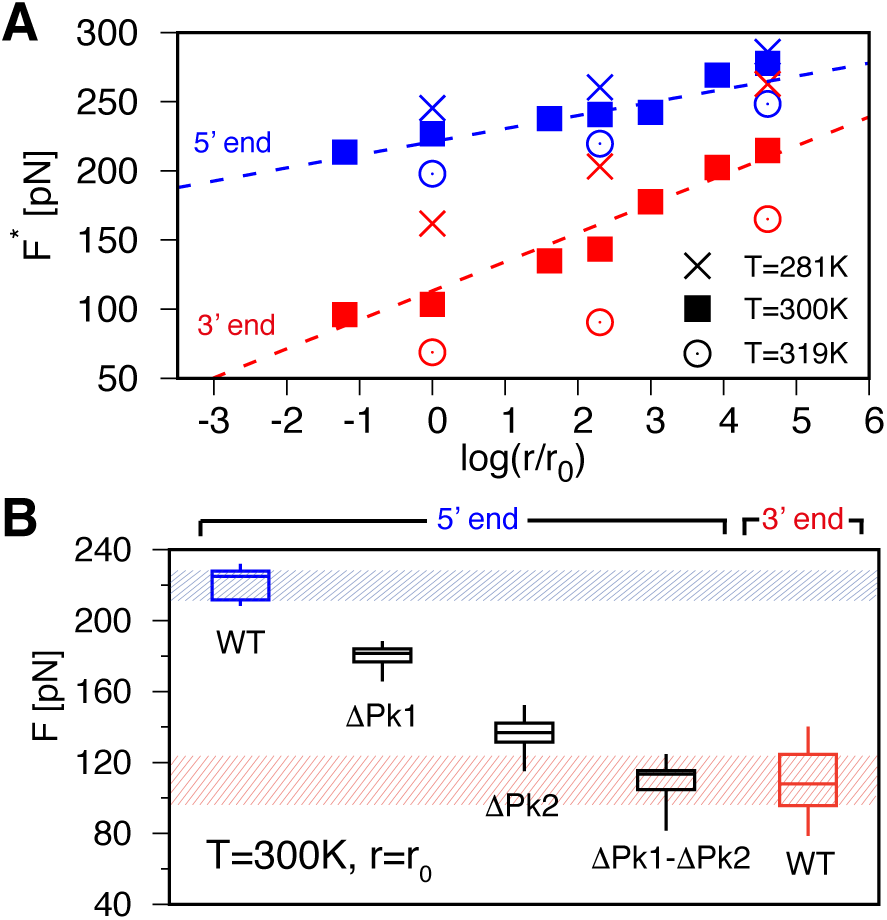
Critical translocation forces for different conditions and variants of Zika xrRNA. (A) Most probable translocation forces from the 5′ and 3′ ends at different temperatures, *T*, and force-ramping rates, *r*; the estimated errors are smaller than the symbols. For clarity only a subset of all considered (*T, r*) combinations are shown, see Fig. S7. The dashed lines show the Bell-Evans relation of Eq. 1 for *T* = 300K with parameters fitted simultaneously on all (*T, r*) datapoints, see main text. (B) Box-whisker plots (centre line, median; box limits, upper and lower quartiles) for the translocation forces at *T* = 300K and *r* = *r*_0_, for the native (WT) xrRNA (5′ and 3′ entries) and for three variants lacking the native contacts in Pk1 and/or Pk2 (5′ entries).

The best-fit parameters are 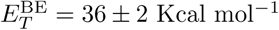 and Δ^BE^ = 4.4 ± 0.4Å for 5′ entries and 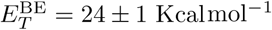 and Δ^BE^ = 2.0 0.1Å for 3′ ones. At room temperature, *T* = 300K these barriers are approximately equal to 60*K*_*B*_ *T* and 41*K*_*B*_ *T* for the 5′ and 3′ ends, respectively.

These barriers describe the system in the absence of a driving force and reflect that translocation is very unlikely to be activated by thermal fluctuations only, especially at the 5′. In the presence of a constant driving force, an estimate of the activation time can be obtained via the so-called Bell-Evans lifetimes [20] and the best-fit parameterization of eq. 1, see Methods. For constant driving forces of 50-100pN, a range accessible to molecular motors[28], the estimated activation time from the 5′ end exceeds by more than two orders of magnitude the one at the 3′ end.

### Directional resistance and stress redistribution

The precipitous unlocking of translocation occurs when the *P* atom at the 5′ end is just about at the *trans* edge of the pore, *z* ∼ − 19.5Å(Supplementary Fig. 3). This triggering event is preceded by a long loading phase during which the driving force, which is exclusively imparted to the *P* atoms inside the pore, propagates an increasing tensile disturbance to the xrRNA that is still on the *cis* side.

The ensuing effects are best discussed by considering how the network of native contacts is distorted during the loading stage. The latter can be quantified in terms of the strain, *s*, of the native interactions for each nucleotide. Negative or positive values of *s* indicate that contacting atoms are respectively drawn closer or further apart than in the unstrained xrRNA structure, see Methods. Kymoplots of the *s* profile, during the loading stage of translocation from both ends, are shown in Fig. 4A and Supplementary Fig. 6.

**FIG. 4.**
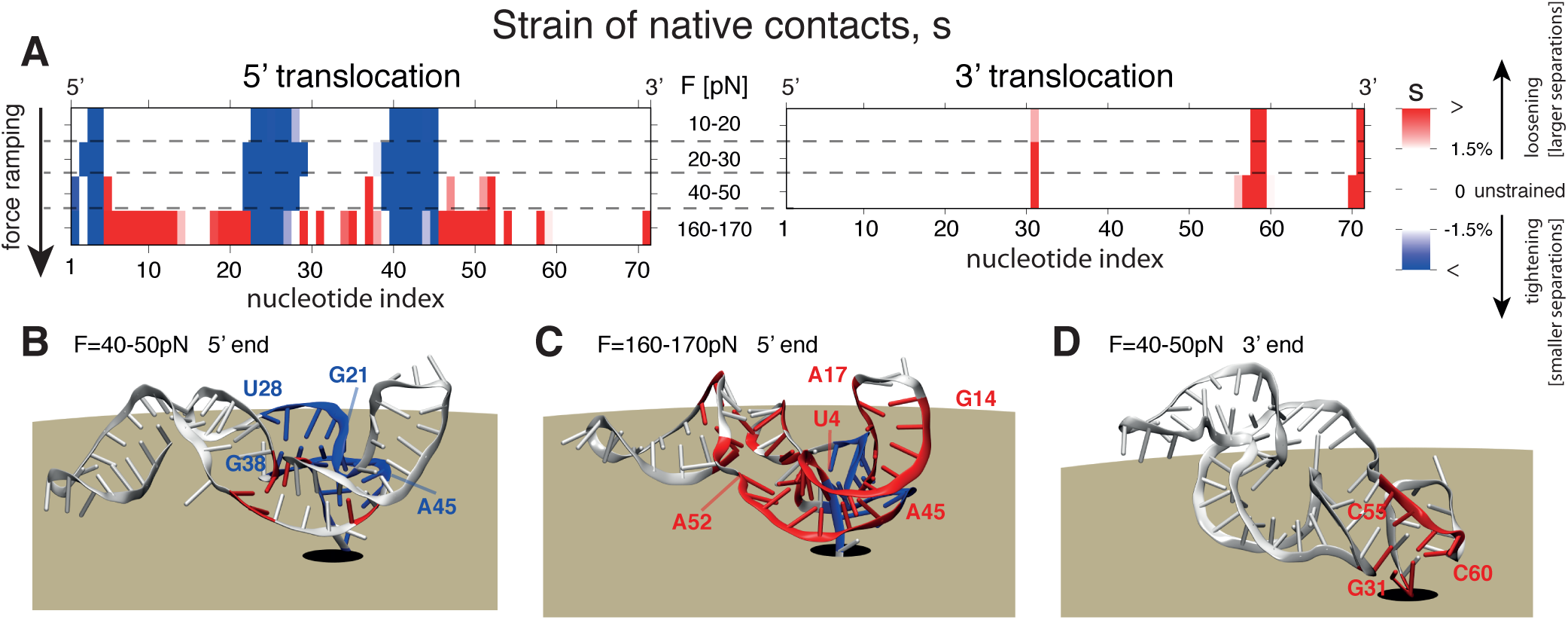
Mechanics of Zika xrRNA directional resistance to translocation. (A) Kymograph of the strain of native interactions, *s*, during the loading stages of force-ramped translocation. Negative or positive value of the strain parameter indicate that natively-interacting atoms are, respectively, closer or further apart than in unstrained xrRNA, reflecting the local tightening or loosening of the structure. The nucleotide-wise profile of the strain was computed for the indicated force ranges over 20 translocation trajectories at *T* = 300K and *r* = *r*_0_, see Methods and Fig. S6. Nucleotides whose native contact network is significant strained (|*s*| *>* −1.5%) are highlighted in panels B-D. The color code is the same as for panel A.

The data and accompanying structures of Fig. 4 clarify that, as the initial transmission of tension along the RNA backbone draws the first nucleotides closer to or into the pore, their network of interacting nucleotides is increasingly distorted. In fact, regions G21-U28 and G38-A45 wrap more tightly around the 5′ end, particularly nucleotide G3 (Fig. 4B). Thanks to this tightening, which has no analog at the 3′ end, the increasing load is distributed over a broad portion of the molecule, which can hence better withstand the dragging force.

Increasing the load at the 5′ end eventually weakens the native interactions in several regions. These include xrRNA portions directly exposed to the traction of the 5′ end (U4-G14) and their partner strands (A17-C22 and C47-A52), see Fig. 4C. Note that the distal region A45-C48 is further disrupted by being pressed against the outer surface of the pore (Fig. 4C and Supplementary Fig. 5). At the 3′ end, instead, no noticeable tightening occurs around the pulled portion. The terminal tract of the P4 helix and their interacting partners in Pk4 and P4 (G31 and C58-C60) are, instead, directly exposed to the unmitigated pulling into the pore. (Fig. 4A,D). The lack of a mechanism for distributing and dissipating stress is what causes the 3′ end, the one processed by replicases, to yield at much lower forces than the 5′ end, the one resisting to exonucleases.

The tightened portion at the 5′ end, highlighted in blue in Fig. 4, is disconnected along the sequence, but is structurally compact. The emergence of a tightened structural element in response to a pulling force, is a distinctive feature of systems with mechanical tensegrity. In these systems, mechanical resistance results from an equilibrium between elements put under traction (positive strain) and other elements that are put under tension (negative strain) by the same applied force. Thus, the 5′ architecture provides a heretofore rare example of intra-molecular tensegrity, where the very pulling of the 5′ end is what boosts resistance to translocation and consequently to degradation.

### Native contacts network and 5′ resistance

We next investigated which set of native contacts and structural elements at the 5′ end are most conducive to the redistribution of tensile stress and translocation resistance. Our model allows for directly pinpointing these critical contributors, namely by switching off selected combinations of native secondary contacts from the interaction potential. In experiments this is not possible, and similar information can be obtained by performing mutation scans where the identity of nucleotides is affected.

Our analysis indicates that the observed resistance is virtually ascribable to the sole contacts in pseudoknots Pk1 and Pk2. These, we recall, include contacts formed by sugar and phosphate moieties, Fig. 1C. The results are illustrated in Fig. 3B as box-whisker plots for the critical translocation forces of the native xrRNA chain, and three variants lacking the native contacts in either of the two pseudoknots, ΔPk1 and ΔPk2, or both, ΔPk1-Pk2.

In variant ΔPk1 the critical force is appreciably lowered, but it still is almost twice as large as for the 3′ end. A stronger variation is seen for ΔPk2, whose critical force is reduced so much that it overlaps with the critical forces for the 3′ end. A complete loss of directional resistance is found for ΔPk1-Pk2, which yields practically the same forces encountered at the 3′ end. This implies a fall-back to a baseline hindrance once Pk1 and Pk2 are disrupted. Of the two sets of pseudoknotted contacts, those in Pk2 are noticeably the most important ones.

### 5′ resistance, free-energy profile and transition state

We finally used the metadynamics framework[29] to characterize 5′ resistance in more detail and beyond the BE analysis of out-of-equilibrium trajectories. In metadynamics, reversible transitions across significant barriers are achieved with an affordable computational expenditure by using history-dependent biases on multiple order parameters at the same time. Building on the centrality of Pk1 and Pk2 contacts on translocation resistance (Fig. 3B) we used the corresponding fraction of native contacts, *Q*_Pk1_ and *Q*_Pk2_, as natural order parameters, together with the amount of pore insertion of the 5′ *P* atom, *z*. To avoid effects related to the refolding of the RNA portion that has passed to the *trans* side, we extended the length of the pore and profiled the free energy up to *z* = −22Å.

A first result of the metadynamics analysis is the dominant translocation pathway shown in Fig. 5A. The pathway projections in the (*Q*_Pk1_, *z*) and (*Q*_Pk2_, *z*) planes clarify that during the initial stages of translocation, *z* from 0 to −10Å, contacts in Pk1 and Pk2 are only modestly disrupted. Further insertion into the pore, *z* from −10Å to −15Å, causes an appreciable loss of Pk2 contacts. The final insertion tract, *z* from −15Å to −19.5Å, produces a concurrent loss of contacts in both Pk1 and Pk2. Representative structures along the pathway to the force-induced transition state are shown next to the pathway, along with two-dimensional projections of the free-energy profile at the same *z* values, Fig. 5B-D.

**FIG. 5.**
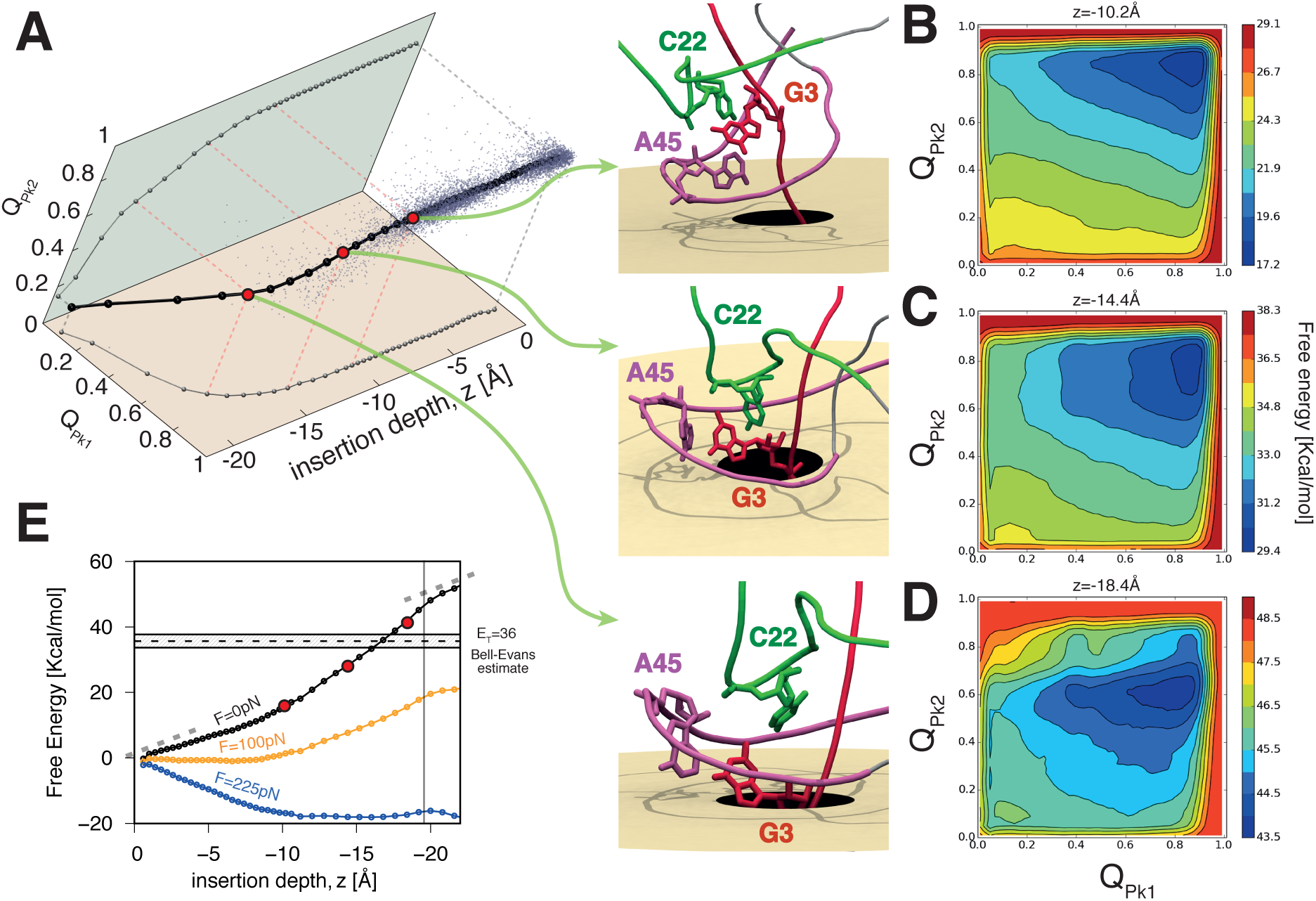
Free-energy profiling of Zika xrRNA translocation. (A) Dominant translocation pathway, from metadynamics analysis, showing the fraction of native contacts, *Q*_Pk1_ and *Q*_Pk2_, versus *z*, the pore insertion depth of the 5′ *P* atom. Its projections in the (*Q*_Pk1_, *z*) and (*Q*_Pk2_, *z*) planes are also shown. The cloud of points shows instantaneous values of the same order parameters in force-ramped translocations at *T* = 300K and *r* = *r*_0_. Representative snapshots at marked values of *z* (red dots) are shown. (B-D) Sections of the three-dimensional metadynamics free-energy at the same marked values of *z*. (E) One-dimensional free-energy profile, obtained from thermodynamic integration and reweighting at the indicated translocation forces. The estimated uncertainty is about 0.55Kcal mol^−1^.

The results clarify that Pk2 interactions yield at *z* ∼ −14Å, when the contacts involving the sugar and phosphate moieties of nucleotides G3-C22 are sheared and disrupted by the drawing of G3 into the pore while C22 is sustained and held in place by the elaborate secondary and tertiary architecture surrounding the 5′ end. The disruption of the non-local contact G3-C22 opens the way for the disruption of other contacts in Pk2 and also Pk1, such as G3 with C44 and A45, for *z* ∼ −18Å.

We note that the dominant pathway established from metadynamics analysis has direct bearings on out-of-equilibrium, force-ramped translocations too. This is clarified by the scattered datapoints in Fig. 5A which represent the same order parameters for conformations sampled from the force-ramped translocation trajectories at *T* = 300*K* and *r* = *r*_0_. The agreement between the dominant pathway and the trajectories has three implications: it corroborates both methods, it indicates that translocation-induced unfolding occurs via a single pathway, and shows that the latter is the same in and out of equilibrium. The results thus cover slower ramping rates than considered in Fig. 3, including those accessible experimentally.

The salient thermodynamics of the process is aptly conveyed by the one-dimensional free energy profile of Fig. 5E, obtained by thermodynamic integration of the metadynamics free-energy at fixed *z*. The free-energy increase up to *z* ≈ −10Å reflects the loss of conformational entropy and native contacts during the loading stage. The steeper rise beyond *z* ≈ −10Å reflects the progressive disruption of Pk2 and Pk1 contacts. The free-energy cost for reaching the depth *z* = −19.5Å, the triggering condition in force-ramped simulations, is about 46Kcal mol^−1^.

Applying constant translocation forces tilts the free-energy profile as shown in Fig. 5E. One notes that at sufficiently high force the slope of the free-energy changes sign at about *z* ∼ −20Å. This value of *z* pleasingly matches the aforementioned triggering insertion depth obtained in force-ramp translocations (Supplementary Fig. 3). The pore insertion depth *z* ∼ −20Å can be interpreted as an effective transition state, embodying the rate-limiting step for force-induced translocations.

In summary, the metadynamics results aptly complement those from force-ramped translocations, giving detailed indications of the free-energy barrier associated with the triggering of translocation from the 5′ end, and of the associated pathway. The combined analysis shows that, in the loading stage, regions G21-U28 and G38-A45, which surround the 5′ end, become more tightly wrapped around it. This mechanism, that has no analog at the 3′ end, stabilizes the network of long-range native contacts around nucleotide G3, hindering their entrance into pore until contacts in Pk1 and Pk2, including G3-C22, are eventually disrupted.

## DISCUSSION AND CONCLUSIONS

Nanopore translocation is a powerful method to probe biopolymers as well as understanding how they interact with processive enzymes [8, 15, 26, 30–35]. Here we used it to clarify the structural determinants of xrRNAs resistance to exonuclease degradation. To this end we carried out hundreds of simulations where an atomistic, native-centric model of Zika xrRNA was translocated through narrow a pore from either end. The setup provides a transparent abstraction of the biological process where the 5′ and 3′ xrRNA termini are engaged and driven through the lumen of enzymes, such as nucleases or polymerases, encountering different directional resistance.

First, we observed that translocation resistance is significantly higher at the 5′ end, the one resisting exonuclease degradation, than at the 3′ one, the one yielding to the unwinding operated by replicases and reverse transcriptases. This holds systematically throughout the many considered combinations of temperatures and force-ramping protocols. It should not go unnoticed that, in our ramping simulations, 3′ -entry translocations not only initiate, but also complete at forces well below those needed to start translocation from the 5′ end. The xrRNA resistance to unfold when pulled through a pore by the 5′ end was also shown to be significantly higher than to unfold by pulling away the two ends (stretching). Bell-Evans analysis indicates that, in the absence of a driving force, the translocation free-energy barriers at the two ends differ by about 12 Kcal mol^−1^. At constant forces of 50-100pN, the estimated difference of activation times exceeds two orders of magnitude, and ought to be detectable experimentally. Incidentally, we note that the directional effect, as well as the different response to translocation and stretching, suggest that experiments where the two ends are pulled apart might lead to results that do not report directly on the resistance offered to electrophoretic translocation and processive molecular motors.

Secondly, the directional effect originates from the different ways in which the pulling effects are propagated from the termini to the rest of the molecule. During the early loading stages, pulling the 5′ end causes regions G21-U28 and G38-A45 to wrap more tightly around it. At later loading stages, interactions between the pulled terminal tract U4-G14 and partner strands A17-C22 and A45-A52 start to yield. At the same time, region A45-C48 also distorts due to the tight contact with the pore surface. Both mechanisms reflect in a compression of the region surrounding the pulled 5′ terminal; this region is compact in space but not in sequence. The tension-induced compression, and the ensuing resistance, make the effect entirely analogous to that of systems with mechanical tensegrity, which oppose tensile forces with compressive ones elicited by the applied tension itself.

As in tensegrity systems, the resistance of the 5′ end is encoded in the architectural organization of the molecule and how it interacts with the pore. By contrast, no such mechanism is available at the 3′ end for its simpler architecture. By selectively removing native secondary interactions, we established that 5′ translocation resistance is primarily encoded in pseudoknot Pk2, and secondarily in Pk1. Once Pk1 and Pk2 contacts are disrupted, no other outstanding barriers are encountered at the 5′ end, and the resistance becomes comparable to that of the 3′ end. The latter fact also rules out the helical asymmetry of nucleic acids, see e.g. refs. [25, 36], as a cause for xrRNAs directional resistance.

Lastly, analysis of free-energy calculations and out-of-equilibrium simulations are consistent at indicating that translocation and the concomitant unfolding from the 5′ end occur via a single dominant pathway. Inspection of the pathway shows that a critical step is the disruption of the interactions of nucleotides G3 and C22, which unlocks the breaking of the other Pk2 and Pk1 contacts eventually triggering the irreversible translocation. Also, comparison with metadynamics results indicates that free-energy barriers estimated with customary Bell-Evans analysis are susceptible to simplifying assumptions on the barrier widths. This finding might be relevant in the analysis of force-ramp experiments too.

In conclusion, these results demonstrate that the structural architecture of xrRNA, which is the sole feature informing our model, is predisposed to generate a profoundly asymmetric resistance to translocation from the two ends. This occurs because mechanical stress is distributed and accumulated in the network of contacts in a way that depends not only on the end that is directly pulled, but also on the regions interacting with it, that become more tightly wound around it. This pulling-induced tightening protects the 5′ end and allows it to better withstand further pulling of the end itself.

Note that this resistance mechanism is altogether different from the catch-bonding behaviour [37] of knotted molecules translocating through narrow pores, which is qualitatively opposite to what observed here. Knotted filaments, particularly those with twist knots, can translocate at sufficiently low forces but jam at high ones[38]. For such systems, jamming happens when the knot, which is pulled against the pore entrance, becomes tight enough that any stochastic sliding motion of the chain along its knotted contour is suppressed [9, 38]. In xrRNA, instead, translocation is stalled at low forces but is unjammed at sufficiently high ones, and moreover in a directional-dependent manner. For xrRNA the initial resistance is offered by the strained network of native interactions that compactify the molecule around the region at the pore entrance, but that eventually yields ad sufficiently high forces.

The results help rationalize and recapitulate in a single structural and quantitative framework not only xrRNA resistance to 5′ degradation from various exonucleases [5], but also the concomitant much lower hindrance to being fully processable from the 3′ in order to be copied by virally encoded RNA-dependent RNA polymerases. Both properties thus find a natural explanation as different manifestations of the same architecture-dependent response. The results also indicate that present-day experimental setups for pore translocation ought to be adequate to capture and reveal the discussed signature effects of the xrRNA directional resistance. In perspective, the structural organization principles that Zika xrRNAs have acquired by evolutionary selection could be co-opted for designing synthetic molecules or meta-materials[39] with directional resistance to translocation and context-dependent tensegrity response.

## METHODS

### Zika xrRNA and system setup

The reference, native conformation of Zika xrRNA was taken from the protein databank [40] entry 5TPY [3]. Simulations of the 71-nucleotide long RNA (1527 heavy atoms) were based on SMOG, a native-centric model retaining the full atomistic structural details [41, 42]. The SMOG potential includes terms for bonded, non-bonded, angular and dihedral interactions. The model is designed for exposing structurally-encoded properties by neglecting effects, such as transient formation of non-native contacts and details of interaction with water and ions. These effects are not expected to be crucial in this specific mechanistic application where, besides, the faster ramping protocols called by explicit solvent treatments would antagonize directional resistance, Fig. 3A. Default values for the relative strength of the potentials and the cut-off contact map were used [42, 43]. The absolute temperature was calibrated similarly to ref. [25] by matching the experimental condition that mechanical unfolding of RNA helices[44, 45] occur at about 15pN at 300K, see Supplementary Methods and Supplementary Table 1.

The *P* atom at the 5′ or 3′ end was primed at the entrance of a cylindrical pore and prevented from retracting from it by a ratchet-like restraint. The pore is 11.7nm-wide and transverses perpendicularly a 19.5Å thick slab, which separates the *cis* and *trans* sides. The pore geometry approximates the lumen of the Xrn1 exoribonucleases, PDB entry 2Y35 [21]. The same SMOG inter-atom excluded volume potential was used for steric interactions of the RNA atoms with the surface of the slab and pore.

### Molecular dynamics simulations

Langevin molecular dynamics (MD) simulations were carried out with the LAMMPS package [46], with proper atomic masses, default values for the friction coefficient and with integration timestep equal to 8.7 × 10^−4^*τ*_*MD*_, where *τ*_*MD*_ is the characteristic MD timescale, see Supplementary Methods. Chain translocation from the *cis* to the *trans* sides are driven by a force, *F*, parallel to the pore axis, and that is exclusively applied to the *P* atoms inside the pore, among which it is equally subdivided at any given time. The driving force is ramped linearly with time *t, F* = *r · t*. In stretching simulations, the 5′ and 3′ *P* atoms are pulled in opposite directions with the same loading protocol. We considered seven equispaced temperature in the range 281-319K, and as many pulling protocols, *r/r*_0_ = 0.3, 1, 5, 10, 20, 50, 100, with *r*_0_ = 6.*×* 2 10^−4^pN*/τ*_*MD*_. Conventional mapping to physical units yields *τ*_*MD*_ ∼ 2.3ps, see Supplementary Methods, which yields *r*_0_ = 270pN/*µ*s, which is conservative for molecular dynamics simulations though still faster than experiments. Note, however, due to the lack of explicit solvent in the model, the effective simulated timescale (pulling rate) reported here represents a lower (upper) bound estimate[47], implying a better agreement with experimental pulling rates. A total of 19 combinations of parameters were explored, see Supplementary Table 2, and for any of them 20 translocation simulations were typically collected for each orientation (5′ → 3′ and *vice versa*). At the reference temperature and ramping rate (*T* = 300K, *r* = *r*_0_) the number of timesteps needed to trigger translocation from the 5′ end is about 7 × 10^8^. Including stretching simulations, more than 1,000 trajectories were run on the Ulysses supercomputer at SISSA.

### Bell Evans analysis

For each orientation, the most probable translocation forces at the 19 combinations of temperature and loading rate, *F* *(*T, r*), were simultaneously fitted with the Bell-Evans expression of eq. 1. The fit consisted of a *χ*^2^ minimization over the free parameters *E*_*T*_, Δ and *v*. *F* *(*T, r*) was established from a Gaussian kernel-density analysis of translocation triggering events, corresponding to a pore insertion depth of the terminal *P* atom equal to z=-19.5Å and −15Å for 5′ and 3′ entries, respectively. The estimated error on *F* *(*T, r*) was set equal to the error of the mean translocation force. The BE waiting time for triggering translocation at a fixed force, *F*, is *τ* = *F*_*β*_*/r*_*F*_, where *F*_*β*_ = *K*_*B*_ *T/*Δ and *r*_*F*_ is the loading rate for which *F* * = *F* [20].

### Strain analysis

We considered the relative strain of a nucleotide, *s* = (*q*^0^ − *q*^*F*^)*/q*^0^. *q*^0^ is the average contact fraction (overlap) of all native interactions of the nucleotide computed at a reference range of low forces, 0-10pN. *q*^*F*^ is analogously computed at force *F*. The contact fraction of two atoms at distance *d* is *q*(*d*) = 1/(1 + [*d/*(1.5 *d*^0^)]^6^), *d*^0^ being their native distance (default cutoff distance of 4Å).

### Metadynamics

Metadynamics simulations [29] were carried out with the COLVAR module [48] using three collective variables: *z, Q*_Pk1_ and *Q*_Pk2_, which are respectively the depth of insertion of the 5′ *P* atom into the pore, and the fraction of native contacts in pseudoknots Pk1 and Pk2. *Q*_Pk_ was computed as 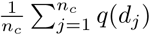, where *j* runs over the *n*_*c*_ native contacts of the pseudoknot. Unbiased simulations, see Supplementary Fig. 8, were used to set the widths of the metadynamics multivariate Gaussians 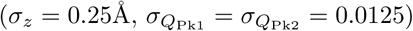, their height (0.15Kcal mol^−1^) and deposition interval (10^3^ timesteps). Retraction of chain from the pore was prevented by a restraining potential, 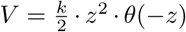, where *k* = 15.6Kcal mol^−1^Å^−2^ and *θ* is the Heaviside function. For the metadynamics analysis, the pore was extended so to reconstructs the free-energy landscape up to *z* = − 22Å, which required runs of 1.5 10^9^ timesteps, Supplementary Fig. 9. A constant force bias of 110pN was used to reduce the time required for filling the basin. The bias was then discounted with a suitable reweighting *a posteriori*. The dominant pathway was obtained by discretizing *z* with intervals of width Δ*z* = 0.8Å and computing the corresponding canonical averages of *Q*_PK1_ and *Q*_PK2_.

## Acknowledgements

This work was partially supported by MIUR, the Italian Ministry of Education. We thank Antonio De Simone for useful discussions.

## Supplementary information

### Supplementary Methods

#### A. Mapping simulation units to physical units

SMOG interaction parameters are in reduced, or simulation units that need to be appropriately mapped or rescaled to match the behaviour of the simulated molecule at the physical temperature of interest.

For our system, which is studied at a single temperature, the mapping of various physical quantities is conveniently expressed in terms of a single rescaling parameter, *α*, as summarised in the table below.

**Supplementary Table I:** Summary of how different physical quantities are mapped from simulation units (LAMMPS implementation of the SMOG model) to actual, physical units. *k*_*B*_ is the Boltzmann constant. Forces in Kcal mol^−1^ Å^−1^ units can be converted to pN by multiplying by 69.5pN mol ÅKcal^−1^.

|  | quantity in simulation units | quantity in physical units |
| --- | --- | --- |
| length | $\sigma^{\text{SMOG}}$ [$\text{\AA}$] | $\sigma = \sigma^{\text{SMOG}}$ [$\text{\AA}$] (no rescaling) |
| mass | $m^{\text{SMOG}}$ [$\text{g mol}^{-1}$] | $m = m^{\text{SMOG}}$ [$\text{g mol}^{-1}$] (no rescaling) |
| Boltzmann constant | $k_B^{\text{SMOG}}$ [$\text{Kcal mol}^{-1} \text{K}^{-1}$] | $k_B = k_B^{\text{SMOG}}$ [$\text{Kcal mol}^{-1} \text{K}^{-1}$] (no rescaling) |
| temperature | $T^{\text{SMOG}}$ [$\text{K}$] | $T = \alpha T^{\text{SMOG}}$ [$\text{K}$] |
| thermal energy | $k_B^{\text{SMOG}} T^{\text{SMOG}}$ [$\text{Kcal mol}^{-1}$] | $k_B T = \alpha k_B^{\text{SMOG}} T^{\text{SMOG}}$ [$\text{Kcal mol}^{-1}$] |
| interaction energies | $\epsilon^{\text{SMOG}}$ [$\text{Kcal mol}^{-1}$] | $\epsilon = \alpha \epsilon^{\text{SMOG}}$ [$\text{Kcal mol}^{-1}$] |
| force | $F^{\text{SMOG}}$ [$\text{Kcal mol}^{-1} \text{\AA}^{-1}$] | $F = \alpha F^{\text{SMOG}}$ [$\text{Kcal mol}^{-1} \text{\AA}^{-1}$] |
| MD timescale | $\tau_{MD}^{\text{SMOG}} = \sigma^{\text{SMOG}} \sqrt{m^{\text{SMOG}} / \epsilon^{\text{SMOG}}}$ [$\text{fs}$] | $\tau_{MD} = \alpha^{-1/2} \tau_{MD}^{\text{SMOG}}$ [$\text{fs}$] |
| force ramping rate | $r^{\text{SMOG}} = \Delta \text{force}^{\text{SMOG}} / \Delta \text{time}^{\text{SMOG}}$ [$\text{Kcal mol}^{-1} \text{\AA}^{-1} \text{fs}^{-1}$] | $r = \alpha^{3/2} r^{\text{SMOG}}$ [$\text{Kcal mol}^{-1} \text{\AA}^{-1} \text{fs}^{-1}$] |

#### B. Temperature-based calibration

For our system, the rescaling parameter *α* can be set by calibrating the mechanical unfolding of helix P4, which is 14-nucleotides long, against the known stretching response of RNA helices.

In stretching experiments RNA helices unfold at critical forces of about *F*_*c*_ = 15pN at ambient temperature, *T* = 300K, equivalent to thermal energies equal to *E*_*t*_ = *K*_*B*_ *T* = 4.1pN nm.

The ratio *E*_*t*_*/F*_*c*_ = 0.273nm defines a lengthscale that can be used to calibrate *α*, since lengthscales are conserved from simulation to physical units.

To this end, we considered various values of *T* ^SMOG^ and, for each of them, simulating the mechanical unfolding of the helix P4 by pulling apart its terminal P’s (atoms indices 1225 and 1505 of PDB entry 5TPY) with a force *F*^*SMOG*^ = *K*_*B*_ *T*^*SMOG*^/0.273nm = (*T* ^SMOG^*/T*)*F*_*c*_ = *α F*_*c*_. We then looked for the temperature at which folded and unfolded states were equally populated at this applied force, as expected at criticality.

As shown in Fig. 1, the matching condition occurred for *α* = *T/T* ^SMOG^ = 3.16. This scaling value is in the same range as those reported in ref. [1], where a similar calibration procedure was used.

Note that, as summarised in Table SI, this mapping also dictates that, like temperature, the reduced energies and forces of the SMOG model must also be multiplied by the same factor *α* to have them in physical units.

Force-ramped translocation and stretching simulation were carried out at various equispaced reduced model temperatures which mapped to the following rounded physical temperature values: *T* = {281, 287, 294, 300, 306, 313, 319}K.

**Supplementary Figure 1:**
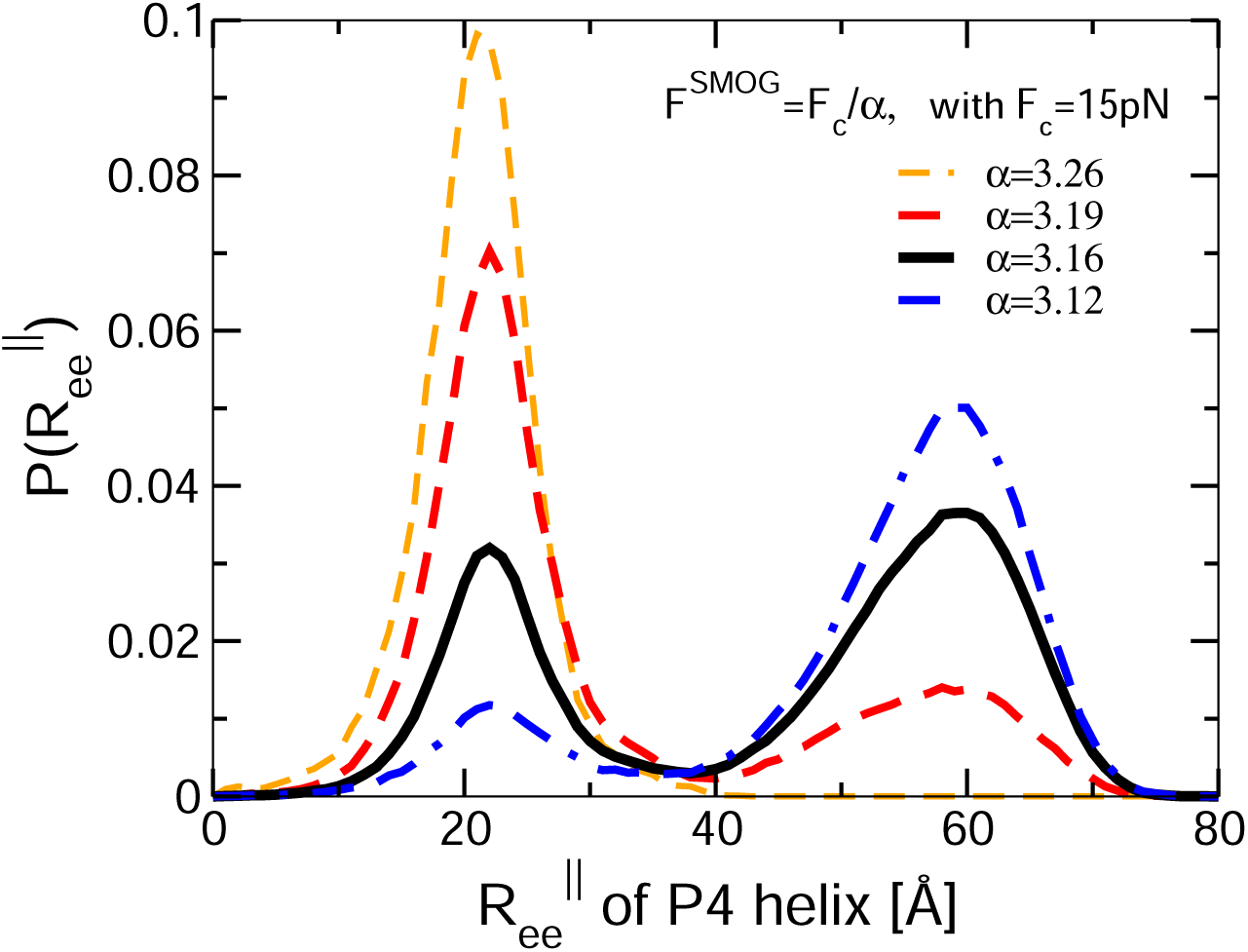
Probability distributions of the longitudinal end-to-end distance of the P4 helix in stretching simulations carried out at the indicated values of *α* = *T/T* ^SMOG^. The stretching simulations were carried out by pulling the terminal P’s of the P4 helix (atoms 1225 and 1505 of PDB entry 5tpy) in opposite directions with a force equal to *F* ^SMOG^ = *α F*_*c*_, where *F*_*c*_ = 15pN, see text. The data were collected over several trajectories, each covering in total from 2.4 × 10^8^ to 5.9 ×10^9^ timesteps at each temperature and force. A balance of folded and unfolded simulations, indicative of the correct combination of temperature and critical force, is observed at *α* = 3.16.

#### C. Mapping of time

A characteristic time for molecular dynamics simulations is given by 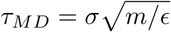, where ^σ^ and *ϵ* are the typical values for the range and strength of the interaction potentials and *m* is the typical mass of the particles. For our model, where we used the actual atomic masses, *m* ∼ 15amu, *σ*^SMOG^ = 4 Å and *ϵ*^SMOG^ = 0.1Kcal mol^−1^, which yields 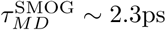.

The Langevin dynamical evolution was integrated with the default integration time step of Δ*t*^SMOG^ = 2fs, corresponding to 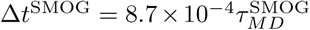 and with inverse damp coefficient equal to 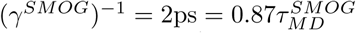.

The MD times in reduced SMOG unit can be transformed to physical units by multiplication by *α*^−1/2^, see Table SI. Accordingly, Δ*t* = *α*^−1/2^ Δ*t*^SMOG^ ≈ 1.1fs and *γ*^−1^ = *α*^−1/2^ (*γ*^*SMOG*^)^−1^ ≈1.1ps. Instead, the force ramping rates in reduced units are transformed to physical units by multiplication by *α*^3/2^, see Table SI.

We recall that mapping of time from reduced to physical units is only approximate since the absence of explicit water molecules in the model might artificially accelerates the dynamics, see e.g. ref. [2].

#### D. Parameter combinations for translocation and stretching simulations

**Supplementary Table II:** List of the 19 combinations of force-ramping rates and temperatures used to profile the translocation response. For each combination we carried out 20 simulations from either of the two ends.

| | $r/r_0$ | $T$ [K] |
| --- | --- | --- |
| 1 | 0.3 | 300 |
| 2 | 1 | 281 |
| 3 | 1 | 300 |
| 4 | 1 | 319 |
| 5 | 5 | 300 |
| 6 | 10 | 281 |
| 7 | 10 | 294 |
| 8 | 10 | 300 |
| 9 | 10 | 306 |
| 10 | 10 | 319 |
| 11 | 20 | 300 |
| 12 | 50 | 300 |
| 13 | 100 | 281 |
| 14 | 100 | 287 |
| 15 | 100 | 294 |
| 16 | 100 | 300 |
| 17 | 100 | 306 |
| 18 | 100 | 313 |
| 19 | 100 | 319 |

**Supplementary Table III:** List of the 11 combinations of force-ramping rates and temperatures used to profile the mechanical stretching response. For each combination we carried out 20 simulations.

| | $r/r_0$ | $T$ [K] |
| --- | --- | --- |
| 1 | 1 | 281 |
| 2 | 1 | 300 |
| 3 | 1 | 319 |
| 4 | 10 | 300 |
| 5 | 100 | 281 |
| 6 | 100 | 287 |
| 7 | 100 | 294 |
| 8 | 100 | 300 |
| 9 | 100 | 306 |
| 10 | 100 | 313 |
| 11 | 100 | 319 |

### Supplementary Discussion

#### E. Translocation and stretching behaviour

**Supplementary Figure 2:**
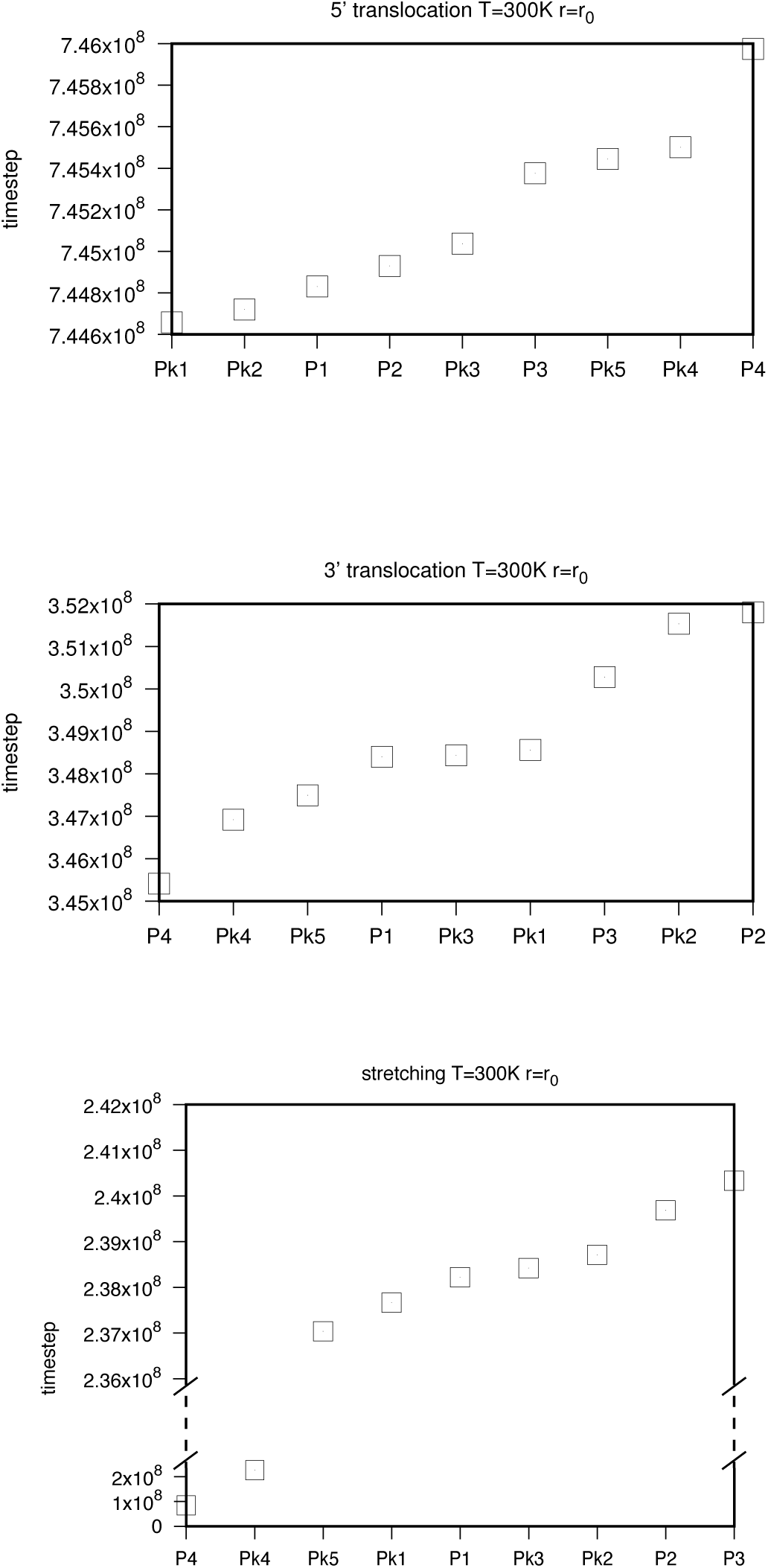
Rupture times of secondary elements during three trajectories: translocations from the 5′ and 3′ ends, and stretching. The rupture event of each secondary element was defined by the fraction of native contacts (overlap) falling below 0.02.

**Supplementary Figure 3:**
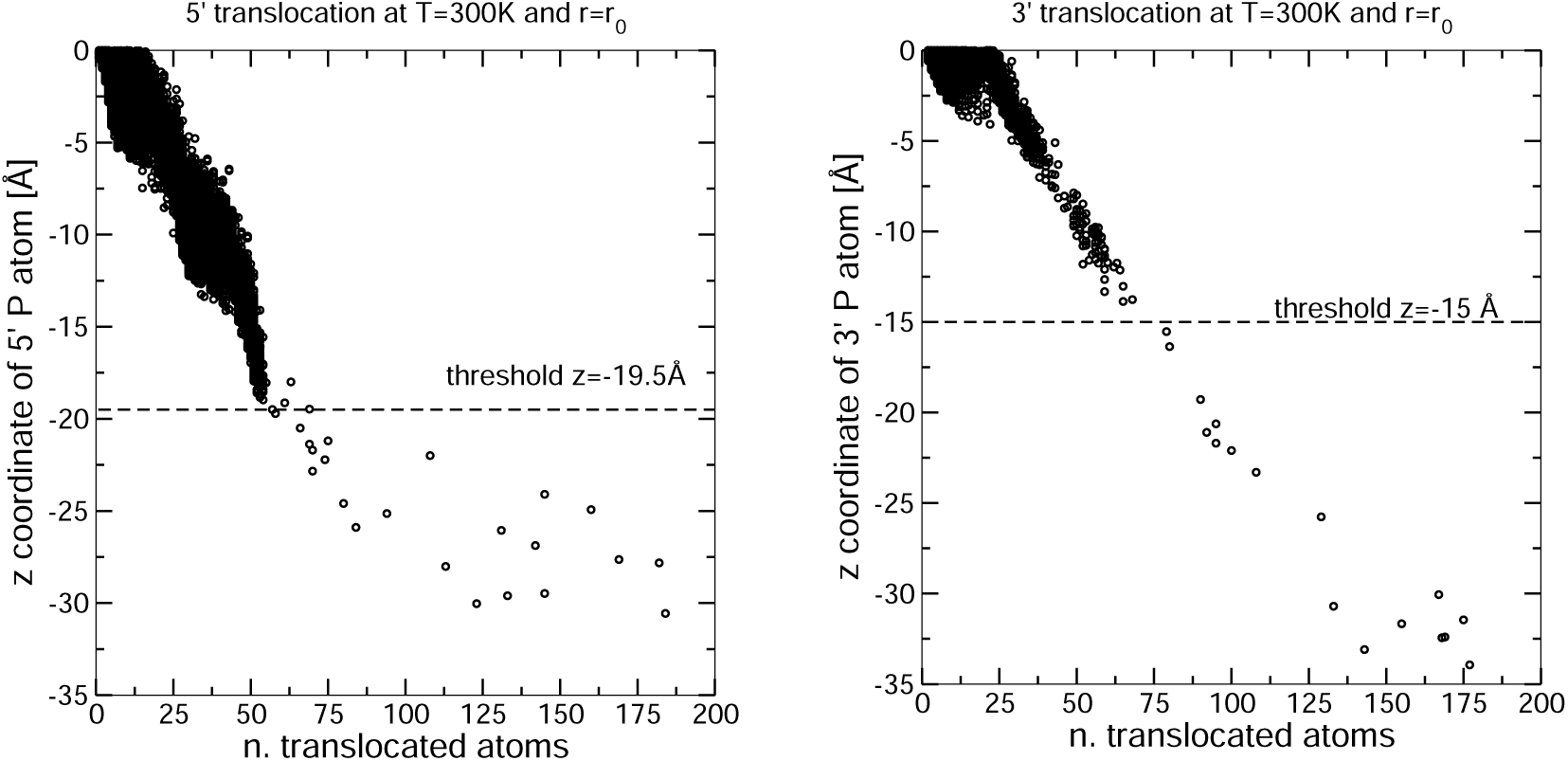
Scatter plot of the *z* coordinate of the *P* atoms at the 5′ and 3′ end versus the number of translocated atoms for force-ramped translocations at *T* = 300K and ramping rate, *r*_0_. In both cases data are for a single run and are, for clarity, limited to the initial stages of translocation. The onset of irreversible translocation is clearly visible as a sparser distribution of the points in the graph. The *z* thresholds defining the triggering of irreversible translocation at the two ends are highlighted.

**Supplementary Figure 4:**
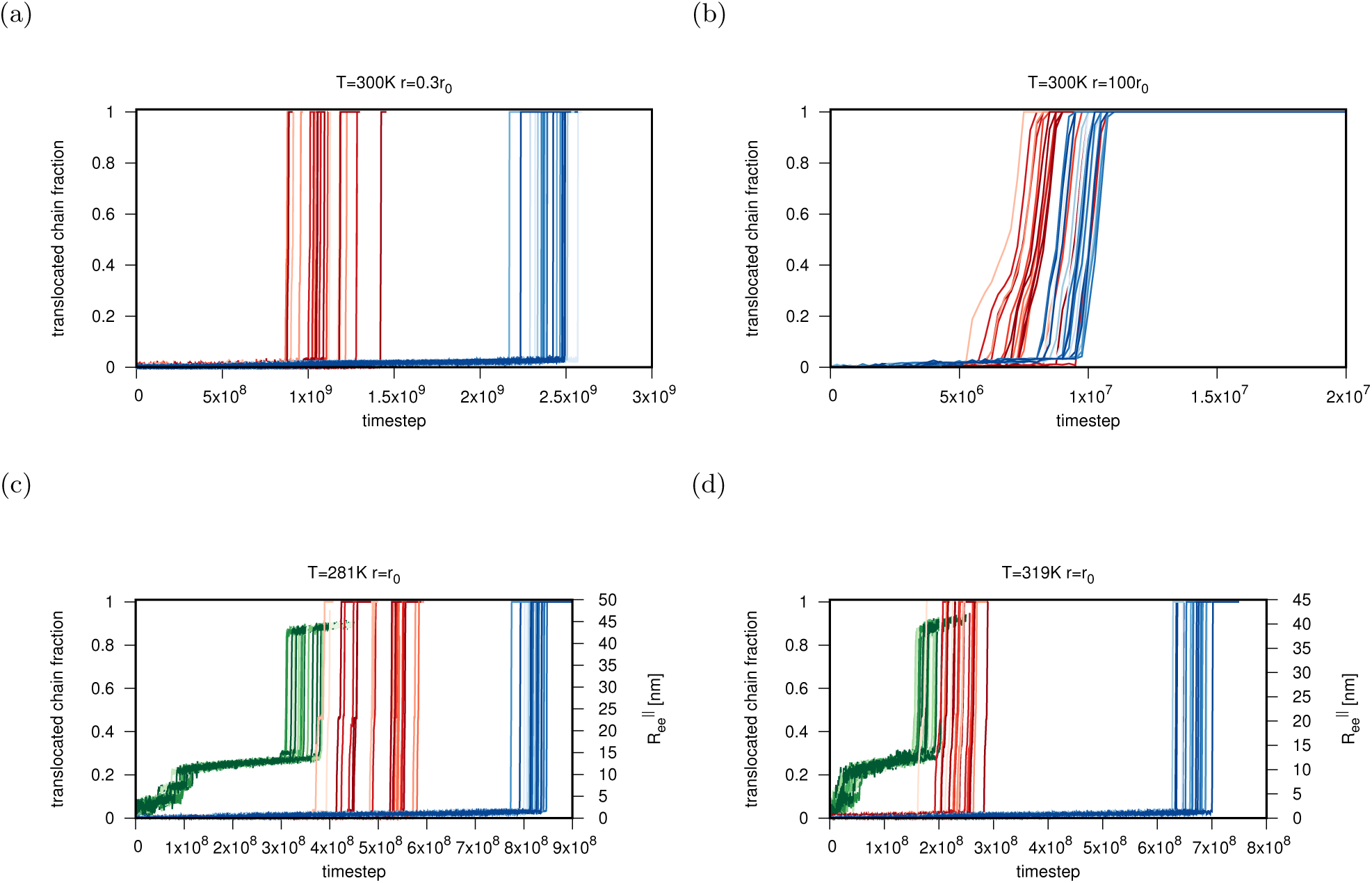
Temporal trace of the translocated fraction of the Zika xrRNA for 5′ (blue curves) and 3′ (red curves) entries for different combinations of temperature, *T*, and ramping rates, *r*. Panels (c) and (d) additionally report the longitudinal end-to-end distance, 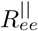 (green curves), as obtained in stretching simulations at the same values of *T* and *r*.

**Supplementary Figure 5:**
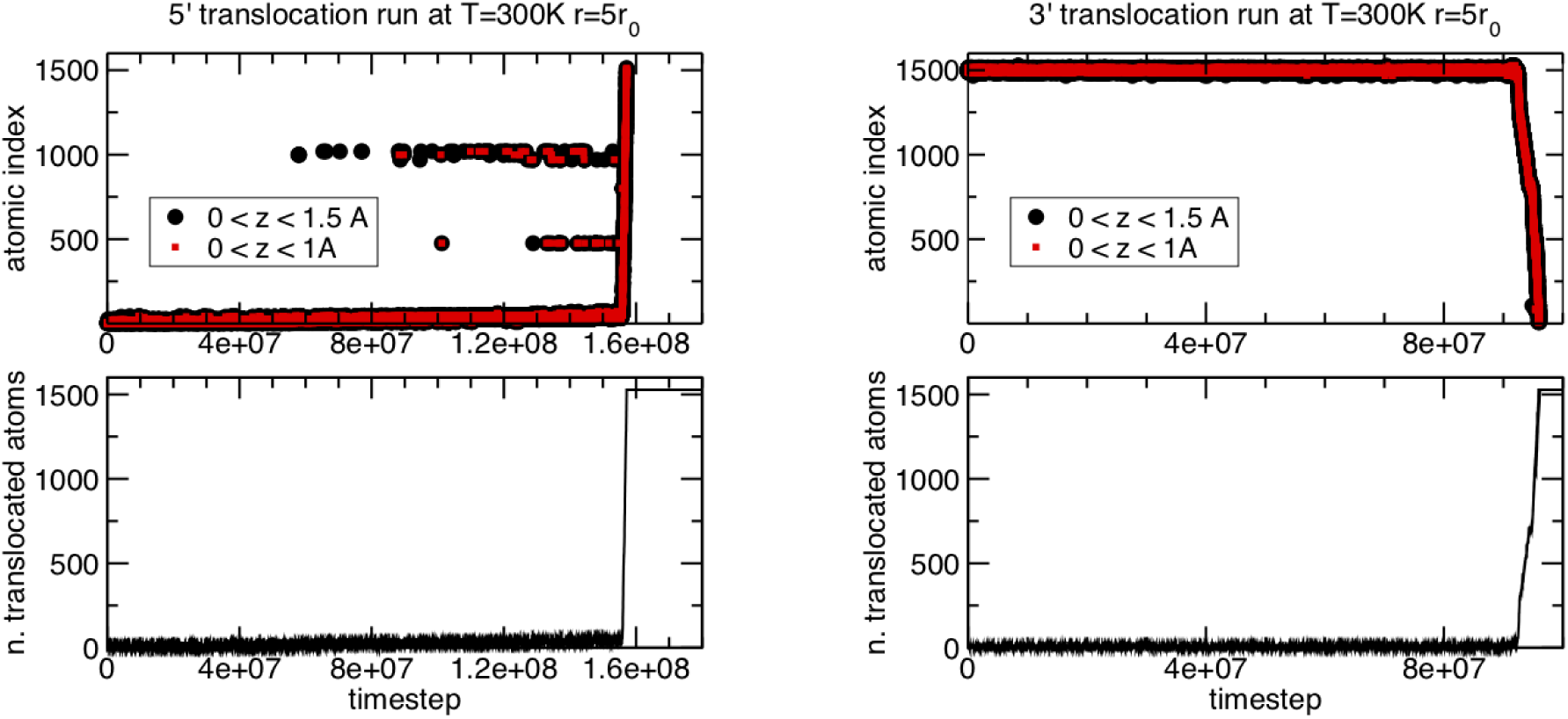
(top) Temporal traces showing the indices of the atoms that during the translocation process have vertical coordinate 0 ≤ *z* ≤ 1.0Åor 0 ≤ *z* ≤ 1.5Å. Data are for a translocation run from either of the two ends at *T* = 300K and *r* = 5 *r*_0_. The two bands visible for 5^*1*^ entries cover the ranges 969-1020 (corresponding to nucleotides A45-C48) and 474-477 (corresponds to nucleotide C22) which are eventually drawn in tight contact with the slab surface prior to the triggering of irreversible translocation. The number of translocated atoms is shown in the bottom panel for reference.

**Supplementary Figure 6:**
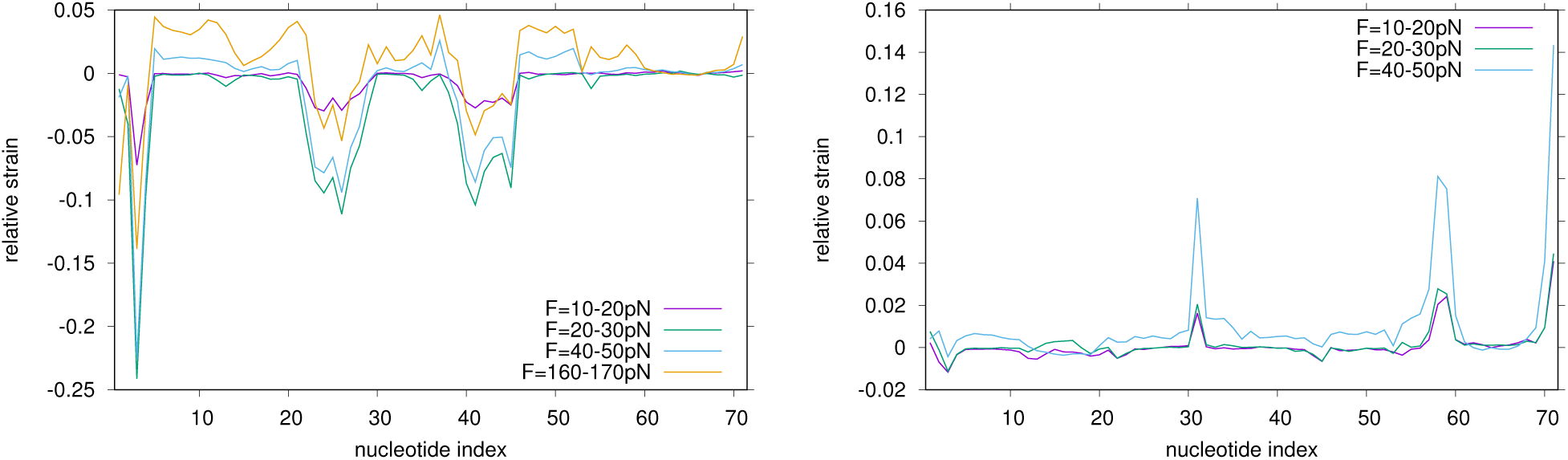
Relative strain profile as a function of the nucleotide index, for translocations from the 5^*1*^ (left panel) and 3^*1*^ (right panel) ends, computed at the indicated force ranges in translocations at *T* = 300K and pulling rate equal to *r*_0_.

**Supplementary Figure 7:**
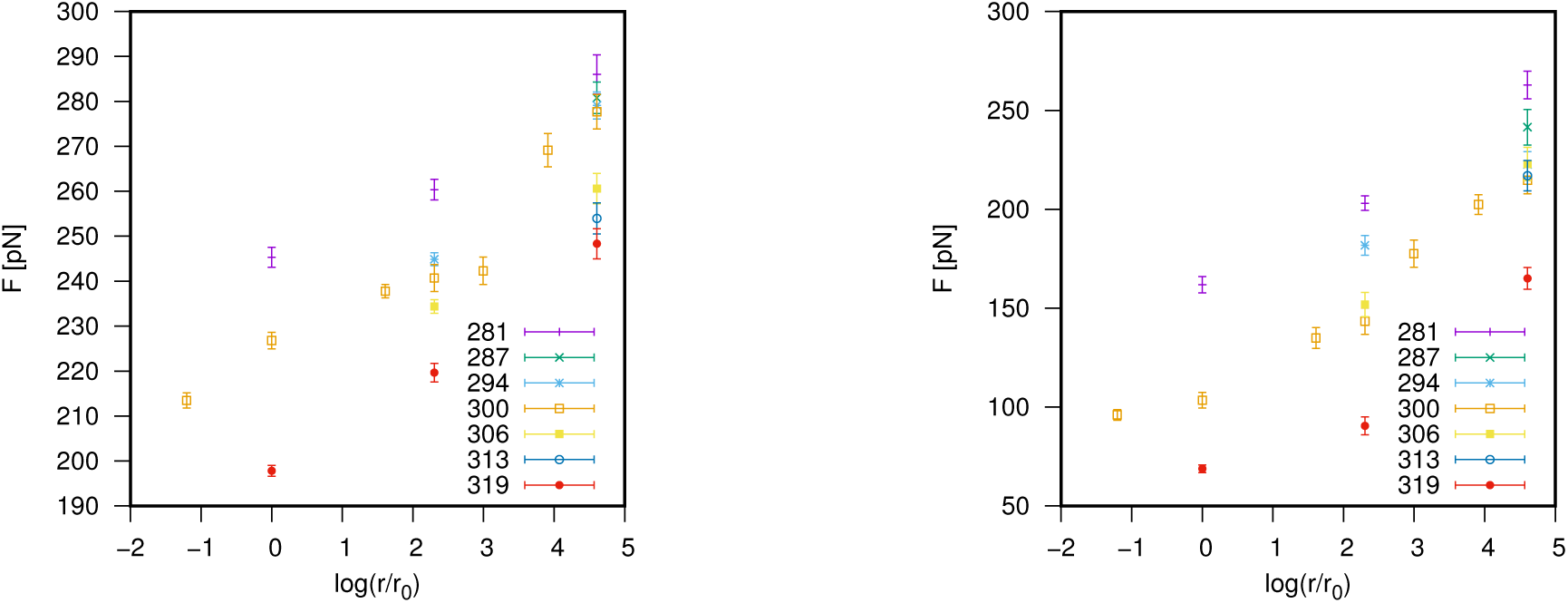
Modal values of the critical (triggering) forces for 5′ (left) and 3′ (right) translocations for the 19 combinations of temperature (different colors) and force ramping rates, *r/r*_0_.

### F. Metadynamics simulations

**Supplementary Figure 8:**
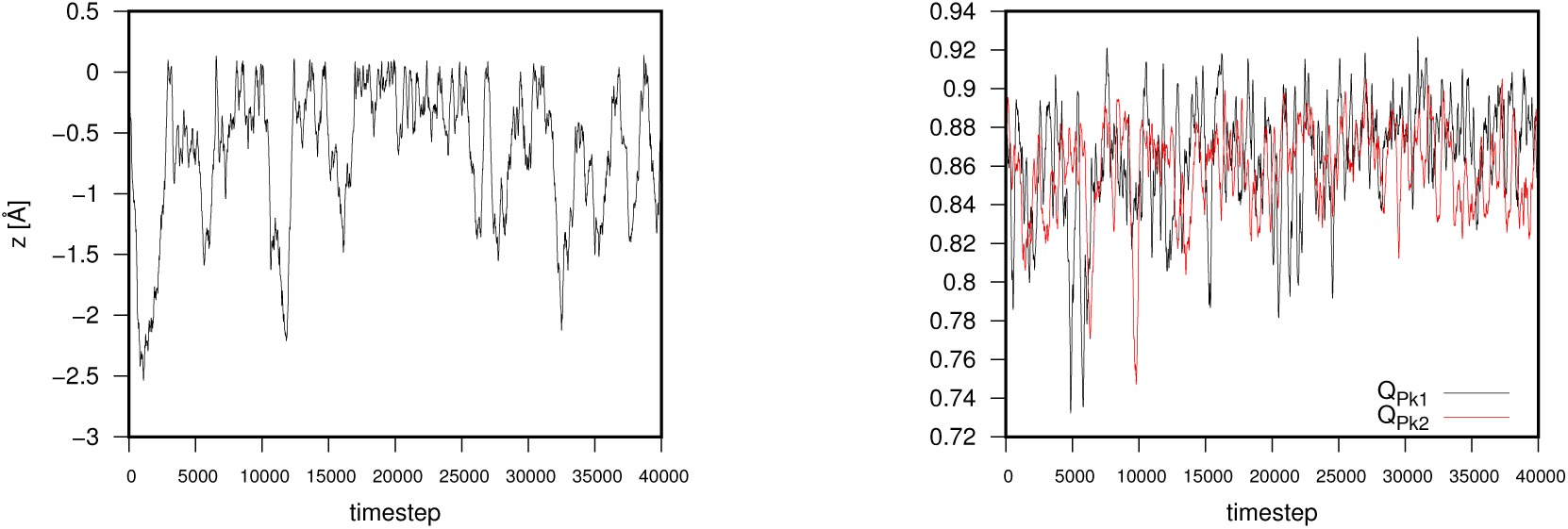
Unbiassed trajectory of the three collective variables used for the metadynamics run. The root mean square fluctuations of the three variables is 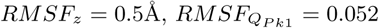 and 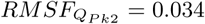. The widths of the multivariate metadynamics Gaussians were set to values smaller than this threshold.

**Supplementary Figure 9:**
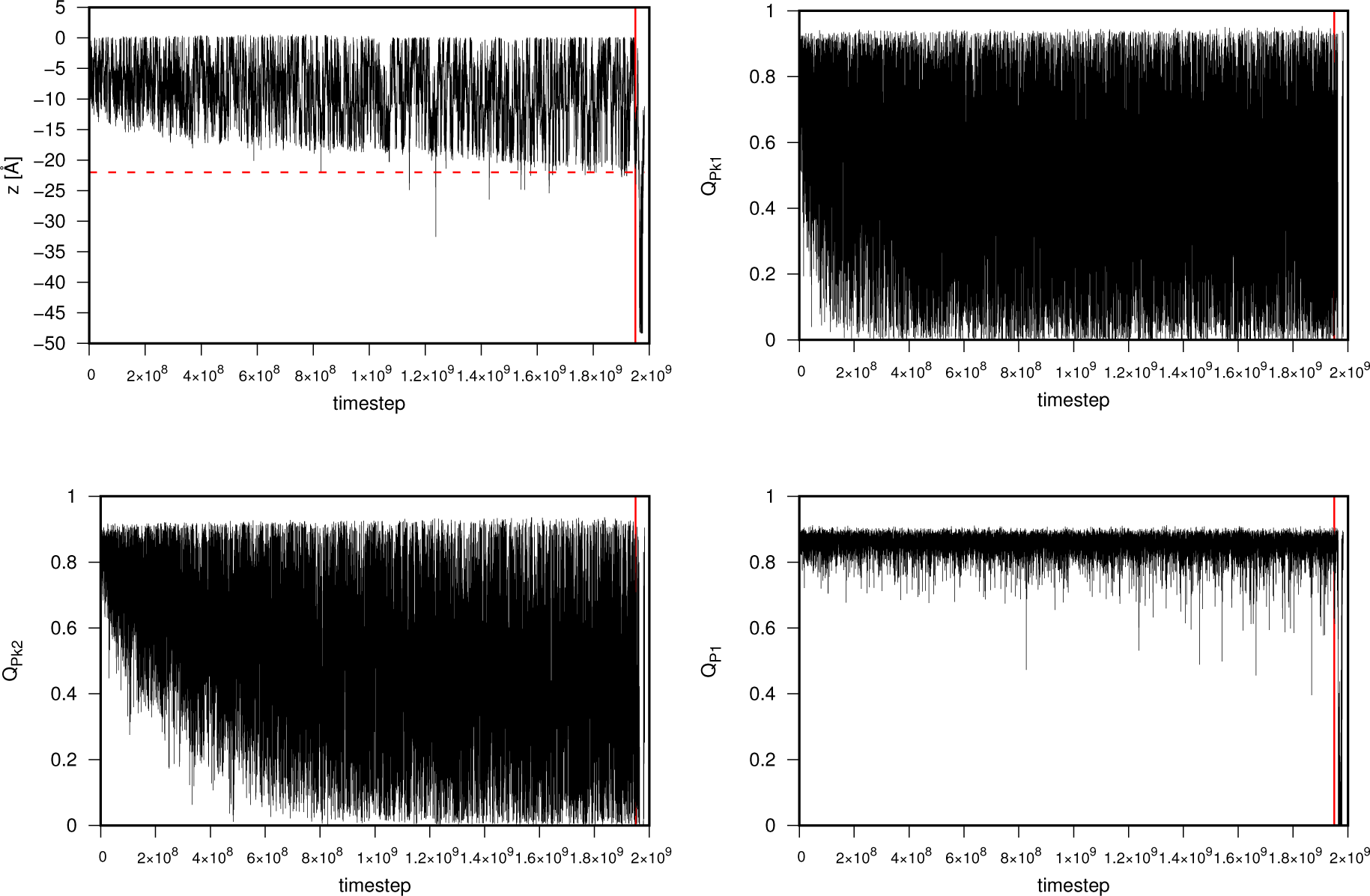
Temporal evolution of the three collective variables *z, Q*_*P k*1_ and *Q*_*P k*2_ during a metadynamics run. The red (dashed) horizontal lines the pore insertion depth *z* = − 22Å. At sufficiently long times, marked by the red (continuous) vertical line, helix P1 eventually breaks and, because its overlap *Q*_*P* 1_ is not included in the collective variables, it is not rapidly reconstituted afterwards. Only the portion of trajectory prior to the breaking of P1 is used to reconstructed the free energy.

**Supplementary Figure 10:**
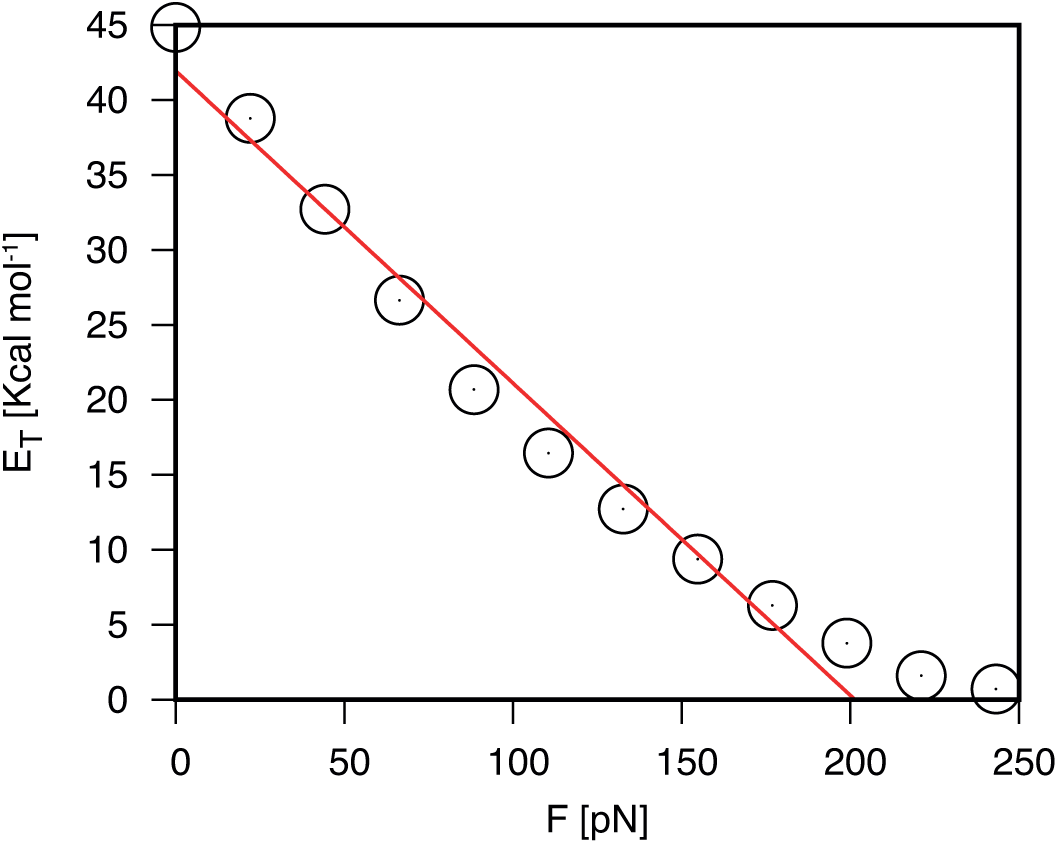
An effective width of the free-energy barrier, can be defined in terms of the variation of the barrier height, *E*_*T*_ upon varying the force, *F*, Δ = − *∂E*_*T*_ */∂F*. The data points show the barrier height for the metadynamics free-energy profile at various values of the added force, *F*. The barrier height, *E*_*T*_, was computed as the difference between the maximum and minimum values of the force-adjusted free energy for −19.5Å ≤ *z* ≤ 0. Linear interpolation of the central datapoints (red line) yields Δ = 13.4Å, comparable to the reduced one of 19.5Å, corresponding to the difference of the *z* values to go from pore entrance to the triggering of translocation at the 5′ end.

